# Using OPM-MEG in contrasting magnetic environments

**DOI:** 10.1101/2021.11.15.468615

**Authors:** Ryan M. Hill, Jasen Devasagayam, Niall Holmes, Elena Boto, Vishal Shah, James Osborne, Kristina Safar, Frank Worcester, Christopher Mariani, Eliot Dawson, David Woolger, Richard Bowtell, Margot J. Taylor, Matthew J. Brookes

## Abstract

Magnetoencephalography (MEG) has been revolutionised in recent years by optically pumped magnetometers (OPMs). “OPM-MEG” offers higher sensitivity, better spatial resolution and lower cost than conventional instrumentation based on superconducting quantum interference devices (SQUIDS). Moreover, OPMs offer the possibility of motion robustness and lifespan compliance, dramatically expanding the range of MEG applications. However, OPM-MEG remains nascent technology; it places stringent requirements on magnetic shielding, and whilst a number of viable systems exist, most are custom made and there have been no cross-site investigations showing the reliability of data. In this paper, we undertake the first cross-site OPM-MEG comparison, using near identical commercial systems scanning the same participant. The two sites are deliberately contrasting, with different magnetic environments: a “green field” campus university site with an OPM-optimised shielded room (low interference) and a city centre hospital site with a “standard” (non-optimised) MSR (high interference). We show that despite a 25-fold difference in background field, and a 30-fold difference in low frequency interference, using dynamic field control and software-based suppression of interference we can generate comparable noise floors at both sites. In human data recorded during a visuo-motor task and a face processing paradigm, we were able to generate similar data, with source localisation showing that brain regions could be pinpointed with just ~10 mm spatial discrepancy and temporal correlations of > 80%. Overall, our study demonstrates that “plug-and-play” OPM-MEG systems exist and can be sited even in challenging magnetic environments.

## 1: INTRODUCTION

Magnetoencephalography (MEG) measures magnetic fields around the head generated by neural current flow (Cohen, 1972). Mathematical modelling of these fields enables generation of 3D images, showing the moment-to-moment evolution of electrophysiological brain activity (Baillet, 2017; Hämäläinen et al., 1993). The fields generated by the brain are small (~10^-13^ T) and to gain sufficient sensitivity, conventional MEG scanners use superconducting quantum interference devices (SQUIDs) which must be cryogenically cooled to liquid helium temperatures (Jaklevic et al., 1964). This places significant limitations on the utility and practicality of the available instrumentation. However, in recent years, MEG system design has been revolutionised by the availability of small, lightweight, and robust optically pumped magnetometers (OPMs) (Alem et al., 2014, 2017; Allred et al., 2002; Borna et al., 2017; Boto et al., 2017; Kominis et al., 2003; Schwindt et al., 2007). OPMs exploit the quantum properties of alkali atoms to measure local magnetic field with high precision. Sensitivity is approaching that of a SQUID, and because the sensors do not require cryogenics, they can be placed closer to the scalp surface, improving sensitivity, spatial resolution, and the uniformity of coverage (Boto et al., 2016; Hill et al., 2020; Iivanainen et al., 2017). Flexible placement of sensors also allows for lifespan compliance (Hill et al., 2019), and assuming background fields are appropriately controlled (Holmes et al., 2018), subjects can move during a scan (Boto et al., 2018). In this way, OPMs are opening new avenues for MEG research, enabling novel experimental design, new subject cohorts, and fundamentally better data. This, coupled with lower purchase and running costs, makes OPMs arguably the most attractive building block for future generations of MEG instrumentation.

Despite the promise, significant hurdles remain for OPMs to overtake SQUIDs as the MEG sensor of choice. Perhaps the biggest barrier relates to the magnetic environment in which systems are housed. Magnetic fields from the brain are much smaller than the fields that exist naturally in the environment. For this reason, all MEG systems are operated inside a magnetically shielded room (MSR), formed from separate layers of high permeability and high conductivity metals (usually mu-metal and aluminium). These act to reduce low frequency, and high frequency interference fields, respectively. However, the requirements for shielding for an OPM system are even more stringent than for SQUID systems; there are three reasons for this: first, OPMs are “zero-field” magnetometers, meaning that their operation is reliant on the background static magnetic field being close to zero (in practice this field can be controlled by “on-board” electromagnetic coils, but the background field must still be < 50 nT). This is distinct from SQUIDs which are relatively unaffected by temporally stationary magnetic fields or “static” field. In most standard magnetically shielded rooms fields, although static field is reduced by flux-shunting in mu-metal walls, the presence of the mu-metal itself leaves a remnant field inside the room, which can be greater than the operational level of an OPM.

Second, once in operation, OPMs have a low dynamic range (~±3.5 nT). This is because as field is increased, the linearity of the OPM response to field is lost; a change in background field of ~3.5 nT would be equivalent to a gain error of 5% (www.quspin.com), raising to 10% for a field change of 5 nT. This means that if the background field drifts over time, or equivalently the OPM array moves with respect to a temporally static field, the OPM measurement will be compromised, and the data rendered useless. Consequently, both low frequency drifts and static field must remain at a level of <3.5 nT (i.e., within 5% gain error) for effective OPM-MEG operation. Third, as in conventional MEG, magnetic interference from the environment degrades signal-to-noise ratio (SNR). However, most OPMs are formulated as magnetometers whereas flux transformers used for conventional MEG are often gradiometers. Magnetometers are more susceptible to magnetic fields from distant sources and so OPM-MEG is ostensibly more susceptible to environmental interference. In sum, the success of OPM-MEG is dependent on extremely accurate control of background fields. This provides a significant challenge, particularly when siting OPM-MEG systems in regions of high magnetic interference (e.g., city centre sites).

In addition to background field, several other challenges exist; for example, minimisation of crosstalk between sensors, optimised array design, robust sensor mounting, and accurate measurement of sensor location and orientation are all requirements for effective OPM-MEG operation. Multiple solutions have been proposed, and a number of effective OPM-MEG arrays are in existence. However, the extent to which one can achieve comparable data from multiple sites – particularly if those sites have different levels of magnetic interference – is unclear. The ultimate success of OPM-MEG will require such cross-site robustness. This, coupled with ease of system use and diminished reliance on an extensive (physics-based) support network, is critical if OPM-MEG is to achieve its full potential and ultimately replace SQUID-based MEG systems.

In this paper, we report the first cross-site OPM-MEG comparison. Specifically, we contrast identical OPM-MEG arrays in very different magnetic environments. The first is a “green field” (campus university) site with an OPM-optimised magnetically shielded room; the second is a city centre hospital site with OPM-MEG installed in an existing (non-optimised) magnetically shielded room. In what follows, we first demonstrate that by a combination of hardware (Holmes et al., 2019) and software (Tierney et al., 2021) approaches for interference reduction, OPMs can be made to work with a similar noise floor in both locations. Following this, at both sites, we capture OPM-MEG data during both a visuo-motor task (well known to generate robust neural oscillatory effects in the beta and gamma bands), and a visual face processing task (known to generate evoked responses from both primary and lateral visual areas) in the same participant. Results from both sites are compared quantitatively, both at the sensor level and following source reconstruction.

## 2: METHODS

All data were collected by the authors. All code for analysis was custom written by the authors using MATLAB.

### 2.1: Site and system descriptions

Our first OPM-MEG system was housed at the Sir Peter Mansfield Imaging Centre, University of Nottingham, UK (SPMIC) – a site with inherently low magnetic interference. The system was housed inside a magnetically shielded room (Magnetic Shields Limited, Kent, UK) comprising 4 layers of mu-metal and a single layer of copper. Static magnetic field inside the room is minimised by a degaussing system (Altarev et al., 2015) which allows demagnetisation of the inner mu-metal walls. Background static field impinging on the array was expected to be ~2 nT, with low frequency (i.e., < 1Hz) drifts in magnetic field of ~0.3 nT, measured over a ten-minute recording (Rea et al., 2021).

Our second site was at the Hospital for Sick Children, Toronto, Canada (SickKids). This is a city centre hospital site with high inherent magnetic interference generated by nearby infrastructure including elevators, a metro-line, parking garages and local construction. The SickKids OPM-MEG system was housed in a MSR (Vacuumschmelze, Hanau, Germany) comprising two layers of mu-metal and a single layer of aluminium (this MSR was previously used for SQUID-MEG). No degaussing was available. The static background magnetic field was expected to be ~30 - 70 nT (~20 times more than the SPMIC site) and maximum field drifts measured over a 10-minute period were expected to be 5-10 nT (~30 times more than SPMIC).

At both sites, the OPM-MEG device was equivalent (Cerca Magnetics Limited, Kent, UK; (Hill et al., 2020)). The array used contained 24 dual-axis zero-field magnetometers manufactured by QuSpin Inc. (Colorado, USA). Each sensor is a self-contained unit, of dimensions 12.4 x 16.6 x 24.4 mm^3^, containing a Rb-87 gas vapour within a glass cell, a laser for optical pumping, and on-board electromagnetic coils for controlling local magnetic field within the cell. Optical pumping polarises the atomic magnetic moments of the atoms in the gas, inducing a bulk magnetisation. In the presence of an external field (i.e., the neuromagnetic field) this magnetisation obeys the Bloch equations and can be exploited to generate a sensitive measure of local field. Two orthogonal components of the local magnetic field (perpendicular to the pumping laser beam) were measured at each OPM *sensor*, giving a 48-*channel* system (note that the OPMs themselves were oriented so field was measured radial to the head, as well as in one tangential orientation). Each channel had an inherent noise floor (environmental interference notwithstanding) of 7 – 10 fT/sqrt(Hz) and a bandwidth of 0 – ~130 Hz. Analogue signals representing the time evolution of measured magnetic fields were fed from the OPM electronics to a National Instruments digital acquisition system (DAQ), via which they were recorded.

Sensors were mounted on the head via a 3D printed helmet (Cerca Magnetics Limited, Kent, UK – Figure 1a). The helmet is made from a lattice which makes it lightweight (700 g) and enables heat to escape from the OPMs (which are heated to an external surface temperature of ~≤40 °C). The lattice also enables free flow of air to the subject’s scalp and contains features for cable management. The helmet contained 64 possible slots for sensor mounting, and the 24 OPMs used were positioned to optimally cover the left parietal and occipital cortices. Figure 1b shows a digital representation of the sensor locations with respect to the brain; the arrows represent the sensitive axes along which field was measured. The coloured surface represents relative sensitivity to dipoles in different brain regions. The left-hand figure shows the array sensitivity to dipoles oriented in theta (i.e., with a polar orientation), and the right-hand figure shows the array sensitivity to dipoles oriented in phi (i.e., with an azimuthal orientation). The colour represents the Frobenius norm of the lead field from each dipole.

**Figure 1:**
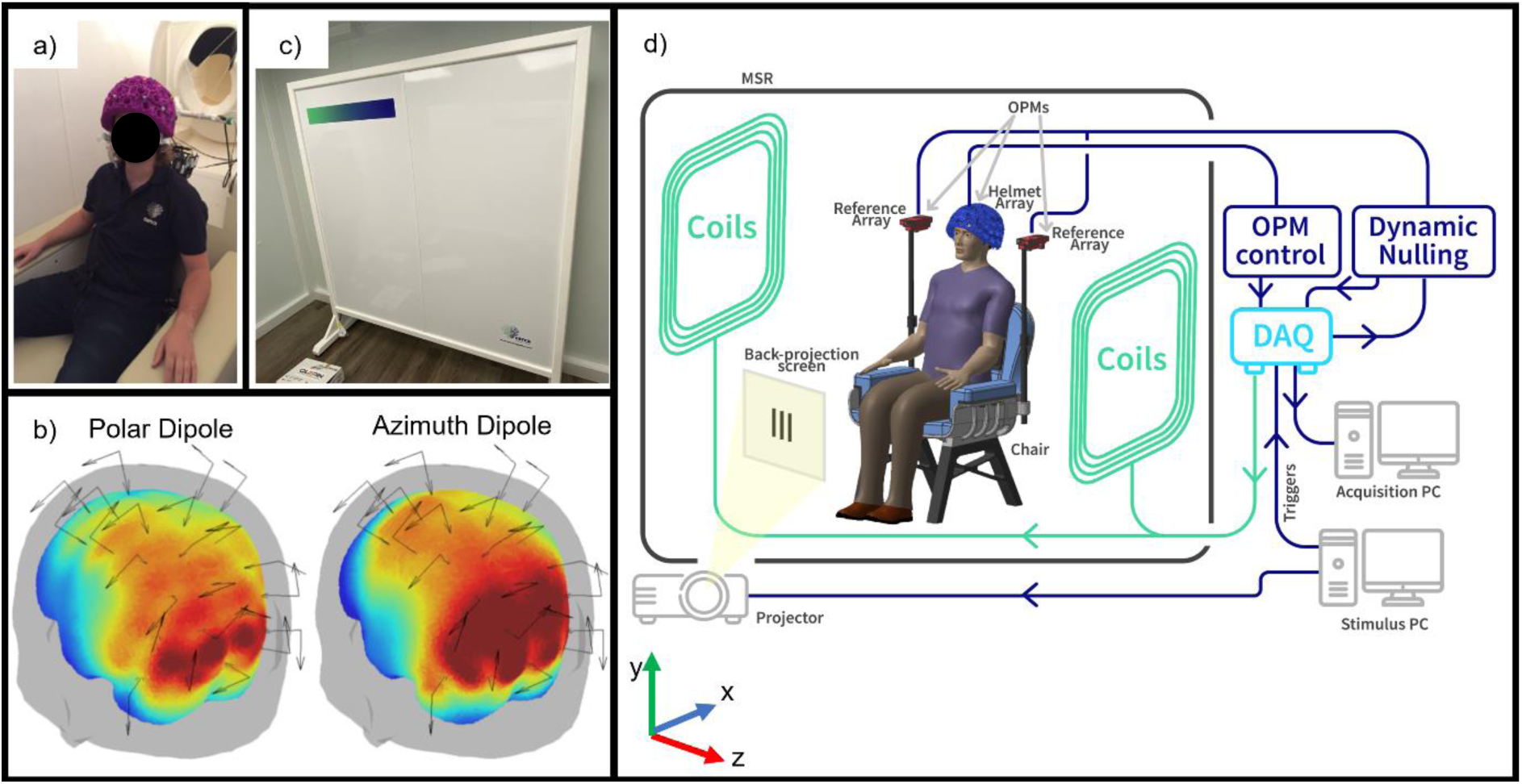
System schematics. a) A lightweight generic helmet designed to fit ~95% of adults. b) OPM placement relative to the head. The coloured surface represents sensitivity to a dipole oriented in the polar (left) or azimuth (right) orientation; we ignore radial dipoles due to the relative insensitivity of MEG to dipoles in this orientation. c) Biplanar coils placed either side of the subject. d) schematic diagram of the Cerca Magnetics OPM-MEG system used at the two sites.

Magnetic field surrounding the OPM helmet was controlled using a set of bi-planar coils placed either side of the participant (Holmes et al., 2018, 2019) (Cerca Magnetics Limited, Kent, UK – Figure 1c). These coils, which are wound on two 1.6-m square planes separated by a 1.5-m gap, generate 3 orthogonal magnetic fields and all 5 independent gradients within a 40-cm cube inside which the participant’s head is positioned. A reference array, placed behind the participant, measures the background field/gradient across the helmet and currents are applied to the bi-planar coils to control this remnant field. At SPMIC, this system of coils was used to remove both the background field and field drift, while at SickKids, only the dynamic changes were cancelled (described below). Consequently, there is a larger static background field at the SickKids site, and so the subject was instructed to it still during acquisition at both sites. Figure 1d shows a schematic diagram of the complete system. Note in addition to the helmet, coils and MSR, a stimulus delivery system was available in both labs to deliver visual stimuli to the participant via back projection through a waveguide in the MSR and onto a screen placed in front of the subject.

### 2.2: Interference rejection methods

As outlined above, the SickKids site was significantly more challenging than the SPMIC site in terms of background magnetic interference. The expected large drifts in background field regularly caused the OPMs to exceed their operational range (which we define at a 5% limit on the accuracy of the OPM gain – which corresponds to a field of ±3.5 nT). Further, we expected the interference inside the room to be significantly worse than that commonly experienced in SPMIC. For this reason, two separate techniques were used to control interference.

- *Dynamic nulling:* To keep the sensors within their operational range of ±3.5 nT, the bi-planar coils either side of the participant were operated in a dynamic proportional-integrative (PI) mode. A complete description of this has been given elsewhere and will not be repeated here (Holmes et al., 2019; Rea et al., 2021). Briefly, a reference array (Figure 1d) consisting of four QuSpin OPMs (two placed either side of the subject’s head, separated by ~30 cm), measured the x, y, and z components of the background field at two locations, as well as the field gradients in the Z direction. The reference magnetometer signals were outputted to a high-speed (60 Hz) PI controller implemented in LabVIEW, which calculates compensation currents which are fed back to the coils. These, in turn, generate temporally changing fields that dynamically compensate <3 Hz changes in the local magnetic field. In this way, we could control both the background field and the drifts inside the MSR.
- *Homogeneous field correction:* To reduce environmental interference after dynamic nulling, we used Homogeneous Field Correction (Tierney et al., 2021). Briefly, the magnetic interference from distant sources, observed by an OPM array distributed over a relatively small volume (i.e., around the head), can be modelled as a spatially homogeneous magnetic field (i.e., we assume that sources of interference are sufficiently distal that the spatial variation of magnetic field over the head volume is negligible). This homogenous field is estimated across the whole array. Then, through knowledge of the sensor orientations, its manifestation at each sensor can be estimated and subtracted from the data. This acts to reduce external interference and improve signal-to-noise ratio. The low rank of the model (i.e., the assumption of field homogeneity) means that there is little risk of removing substantial neural signal, which has marked spatial variation across the array.

### 2.3: Data collection

To test the effects of interference rejection, 5 minutes of empty room data were recorded at each site, with and without dynamic nulling. The data with dynamic nulling were further processed using homogeneous field correction. In all three cases (i.e., no correction, dynamic nulling, and dynamic nulling + homogeneous field correction) the noise floor was assessed quantitatively. The noise data were processed by calculation of power spectral density using Welch’s periodogram method, to give an accurate representation of the noise floor as a function of frequency.

Following empty room recordings, we acquired human MEG data in a single subject. The participant was a male, aged 26, right-handed. We performed two experimental paradigms, both well known to produce robust neuromagnetic effects. The first task was a *visuo-motor* paradigm (Hoogenboom et al., 2006; livanainen et al., 2020). Each trial comprised 1 s of baseline measurement followed by visual stimulation in the form of a centrally presented, inwardly moving, maximum-contrast circular grating. The grating subtended a visual angle of 7.6 degrees at both sites, and was displayed for a jittered duration of either 1.6 s, 1.7 s or 1.9 s. Each trial ended with a 3-s baseline period, and a total of 100 trials was used. During baseline periods, a fixation dot was shown on the centre of the screen. The participant was instructed to perform abductions of their right index finger for the duration that the stimulus was on the screen to ‘activate’ primary motor cortex. We expected to measure simultaneous fluctuations of beta oscillations in motor cortex, and gamma oscillations in visual cortex. The second task was a *face processing* paradigm. Here, the participant was asked to passively view a series of images, each containing a face. In a single trial, a face was displayed on a screen for 300 ms; this was followed by a rest period of jittered duration (1900 ± 181 ms) during which a fixation cross was shown. A total of 100 trials was recorded. This task is well known to generate robust evoked responses both in primary visual cortex (at a latency of ~100 ms) as well as the fusiform area (at a latency of ~170 ms) (Bentin et al., 1996; Halgren, 2000; Taylor et al., 2001). For both paradigms the subject was seated and free to move but not encouraged to do so. MEG data were acquired at a sample rate of 1200 Hz. Each paradigm was independently run 5 times in the same subject at each of the two locations (SPMIC and SickKids).

Following data collection, a 3D optical camera was used to generate a digital model of the location of the helmet (and thus the sensors) relative to the brain anatomy (Hill et al., 2020). A digitisation of the participant wearing the helmet was acquired using a Structure Core 3D scanner (Occipital Inc., San Francisco, CA, USA). This was followed by a second digitisation with the helmet removed and the participant’s hair tied back (to smooth the digitisation of the top of the head). Finally, a structural MRI of the participant’s head was acquired (using a 3T Phillips Ingenia MRI scanner running an MPRAGE sequence with a spatial resolution of 1 mm). An electronic Computer-Aided Design (CAD) file of the helmet with the exact locations and orientations of the sensors was aligned to the first digitisation (of the helmet relative to the face) using 9 identifiable reference points on the helmet. The first digitisation was then aligned to the second using identifiable facial features (e.g., the nasion, the alar facial groove either side of the nose, cheek bones) and an iterative closest point (ICP) algorithm used to fine-tune this alignment (implemented in MeshLab (Cignoni et al., 2008)). The second digitisation was then aligned (using the same method) to the head/face surface extracted from the MRI. This procedure allowed a complete co-registration of the sensor locations and orientations to the anatomical MRI. This would be used later for modelling source locations.

### 2.4: Data analysis

For each recording, following homogeneous field correction, data were bandpass filtered (between 1 and 150 Hz for the visuo-motor paradigm, and 2 and 40 Hz for the face processing paradigm). Bad trials, defined as those in which the standard deviation of the signal at any one sensor was greater than 3 times the average standard deviation of the signal at that sensor across all trials, were removed. Visual inspection of the data confirmed this simple algorithm was successful at removing trials with excessive noise. Following this, we analysed data first in sensor space, and then via source modelling:

#### Sensor space visualisation

For the *visuo-motor task*, data were further filtered into the beta (13 – 30 Hz) and gamma (35 – 60 Hz) bands. A Hilbert transformation was applied to these filtered data, with the absolute value of the resulting analytic signal being used to generate an amplitude envelope (Hilbert envelope) showing modulation of oscillatory amplitude in each band. The envelope was averaged across trials and baseline corrected (the baseline was calculated over the −3.4 s < t < −2.5 s time window, relative to stimulus offset at t = 0 s). The average envelopes for all 5 experimental runs at each of the two sites were then averaged and the standard deviation between runs was found to assess repeatability. This procedure was run for every channel.

To assess sensor space field topography of the beta and gamma band signals, we computed signal to noise ratio (SNR) at each channel. The trial averaged envelope was divided into an “On” window (i.e., when the stimulation was on; −2 s < t < −0.5 s) and an “Off” window (i.e., when the stimulus was off; 0.5 s < t < 2 s). The SNR in the gamma-band was calculated as the difference in mean signal between the windows, divided by the standard deviation of the signal in the Off window. Similarly in the beta-band, the SNR was calculated as the difference in signal means between the two windows, divided by the standard deviation in the On window (note this was to avoid misrepresentation of SNR due to the beta rebound; note also, since the beta amplitude was expected to decrease during stimulation, beta band SNR was expected to be negative). The resulting SNR values were plotted as a flattened topographical map, across sensor locations, to visualise the sensor-space topography of the beta and gamma-band responses. Two separate topographies were derived, one for the radially oriented field, and one for the tangentially oriented field. A time-frequency spectrum (TFS), alongside averaged envelopes for beta and gamma bands, were also constructed for the channels with the highest SNR. The TFS was derived by sequentially filtering signals into overlapping bands, computing the envelope of oscillatory power, averaging over trials, and concatenating in the frequency dimension.

For the face processing task, trials were averaged and baseline corrected (with baseline calculated in the 1 s < t < 2 s time window; t = 0 s corresponds to onset of the face stimulus). The trial-average response for all 5 runs at each site were averaged and the standard deviation found to assess repeatability. The “best” sensor (i.e., the sensor that showed the largest response) was assessed by measuring the range of the trial-averaged signal in the 0.1 s < t < 0.2 s window. A field map was produced showing the field topography at the time of the largest peak in the evoked response. Again, field maps were made for radially and tangentially oriented fields.

#### Source modelling

For both paradigms, source modelling was performed using a vector beamformer. The brain was divided into 4-mm cubic voxels, and at each voxel location, beamformer reconstructed source estimates were made for sources in the polar and azimuth orientations. To generate a visualisation of task induced signal modulation across the brain, a pseudo-t-statistical approach was used to contrast source power in active and control windows. For both tasks, the forward solution was calculated assuming a dipolar source, and a single-shell uniform volume conductor head model (Nolte, 2003) created using FieldTrip (Oostenveld et al., 2011).

For the visuo-motor task, the active and control windows were −1.5 s < t < −0.5 s and 0.5 s < t < 1.5 s (relative to stimulus offset) respectively. Images showing the spatial signature of modulation in oscillatory power were generated for both the beta and gamma bands. Beamformer weights were calculated independently for each band, with the covariance matrices generated using a time window spanning the entire experiment. The covariance matrices were regularised using the Tikhonov method with a regularisation parameter equal to 5% of the maximum eigenvalue of the unregularized matrix. Based on the pseudo-t-statistical images, a peak location showing maximum oscillatory power modulation was determined, and a signal from this location extracted, again using a beamformer. Here, data covariance was calculated in the 1–150 Hz band and beamformer weights were used to generate a ‘virtual sensor’ time course. Note that a single dipole orientation was chosen to maximise the signal to noise ratio at that location. A time-frequency spectrum was then constructed and averaged over all 5 runs for each site.

For the face processing task, the active and control windows were 0.075 s < t < 0.175 s and 1.075 s < t < 1.175 s relative to stimulus onset respectively. Images showing the spatial signature of modulation in task evoked power were generated. The covariance matrix was generated using data filtered in the 2 – 40 Hz and a time window spanning the entire experiment. Again 5% regularisation was used. Two dipole locations were selected – one in the primary visual cortex (MNI coordinates: (−8, −100, 7) mm), and the other at the peak of the average T-stat for each site (in the left fusiform gyrus for both; MNI coordinates: (−45, −60, −10) mm) – and a signal in each location was reconstructed using the beamformer. Evoked responses were generated by averaging over trials. These responses were then averaged across all 5 runs for each of the two experimental sites, and a standard deviation calculated to assess robustness.

## 3: RESULTS

### 3.1 Rejection of interference

Figures 2a and 2b (left panels) show time-series data for a representative sensor placed in an empty helmet at the central region of the bi-planar coils, recorded over 5 minutes. The two black dashed lines at ±3.5 nT represent a field change corresponding to a 5% change in sensor gain (we would deem sensors inoperable at fields outside of this range). The right-hand panels of Figures 2a and 2b show the mean power spectral density over all 24 sensors placed in the helmet, with the inset axes showing data at frequencies <5 Hz. Here, the black dashed line is at 15 fT/sqrt(Hz); in the absence of external interference (i.e., considering only noise inherent to the sensors) we would expect the power spectral density to be below this line at frequencies above ~5 Hz (most OPMs have inherent noise of between 7 – 10 fT/sqrt(Hz)). For this reason, we deem 15 fT/sqrt(Hz) as the ‘target’ baseline noise level (at which inherent sensor noise dominates). In the plots, the blue lines show raw data (i.e., with no dynamic nulling or mean field correction); the red lines show data with dynamic nulling, and the yellow lines show data with both dynamic nulling and homogenous field correction (HFC) applied. Figure 2a shows the case for SickKids; Figure 2b shows the case for SPMIC.

**Figure 2:**
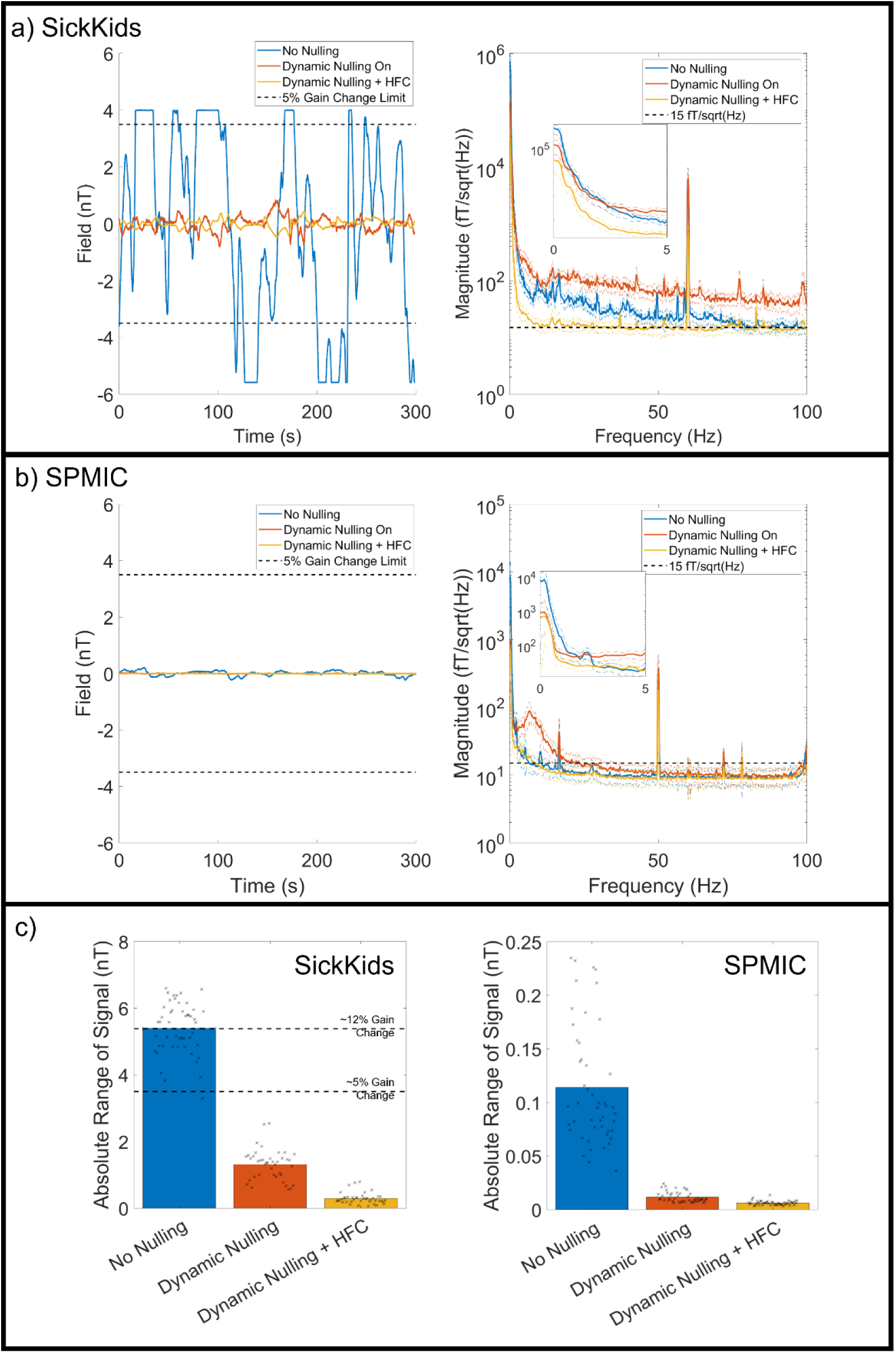
Interference rejection. a) SickKids empty room recording. Raw data for a single representative sensor are shown on the left for the No Nulling Recording (blue), Dynamic Nulling Recording (red), and the Dynamic Nulling Recording with Homogenous Field Correction (HFC) applied (yellow). A dashed line at 3.5 nT represents a gain change in the signal of 5%; if field increases above this line the sensor is non-operational. On the right, the power spectral density (PSD) of each recording is shown, with the inset showing the differences at low (<5 Hz) frequencies. b) Identical to a) for the SPMIC site. c) Left: the average absolute range (i.e., the largest change from zero) for each recording for the SickKids site. The black dashed lines show the 5% and 12% gain change limits. In both plots, the black crosses show the values for each channel. Right: The same plot for the SPMIC site.

At the SickKids site, when no nulling is applied, the background field drifts cause the sensor to regularly exceed its operational range. When dynamic nulling is applied, the sensor is kept well within its operational range, but the noise floor above 3 Hz is raised. HFC removes the majority of the interference, bringing the noise floor close to 15 fT/sqrt(Hz). At the SPMIC site, even with no nulling the sensor is well within its operational range. Dynamic nulling reduces the amplitude of the low frequency interference but again increases interference above 3 Hz. HFC again corrects the noise floor above 3 Hz to a level similar to the no nulling case.

These example results are formalised in Figure 2c. Here, the left panel shows the mean absolute range (i.e., the absolute value of the maximum change from zero) for all 48 channels in the SickKids array. The black crosses represent the individual values for each channel. In the no nulling case, the average range is in excess of 5 nT, which corresponds to a gain change in excess of >10%, and all but one channel exceed their operational range at some point during the 5-minute recording. However, when dynamic nulling is applied, all channels remain within their operational range, and HFC reduces this further. The right panel shows the equivalent data for the SPMIC site (note the difference in the y-axis scale).

These data show clearly that dynamic nulling can be used to maintain sensor operation, even at a site where there are large changes in background field. However, this comes at the cost of increases in higher frequency interference which is generated by noise in the coil current drivers. Consequently, with only dynamic nulling, the background noise is above the 15 fT/sqrt(Hz) target. However, HFC corrects this, as well as supressing other background interference. On average in the 10-Hz to 40-Hz frequency range, following both dynamic nulling and HFC, the measured noise floor across the array (mean ± standard deviation) was 16.4 ± 2.8 fT/sqrt(Hz), and 10.4 ± 2.1 fT/sqrt(Hz), for the SickKids and SPMIC sites respectively.

### 3.2 Data rejection

In the human experiments, we rejected trials with high levels of interference. These data, for both sites, are shown in Table 1. At the SickKids site, on average 22% of the trials for the visuo-motor task had to be discarded (~2 minutes of data) due to unpredictable interference; likewise, 20% of the trials were discarded for the face processing task. At the Nottingham site, these values were reduced significantly for the face processing task (5.6% of trials), while only slightly for the visuo-motor task (16% of trials). These data will be further discussed in Section 4.

**Table 1:** Trials rejected. The number of trials (out of 100) rejected for each run at each site.

| VISUO-MOTOR TASK | SPMIC<br>NUMBER OF BAD TRIALS<br>(OUT OF 100) | SICKKIDS<br>NUMBER OF BAD TRIALS<br>(OUT OF 100) |
| --- | --- | --- |
| RUN 1 | 12 | 10 |
| RUN 2 | 13 | 12 |
| RUN 3 | 26 | 12 |
| RUN 4 | 13 | 28 |
| RUN 5 | 18 | 50 |
| FACE PROCESSING TASK |  |  |
| RUN 1 | 2 | 4 |
| RUN 2 | 12 | 29 |
| RUN 3 | 3 | 27 |
| RUN 4 | 4 | 22 |
| RUN 5 | 7 | 20 |

### 3.3 Visuo-motor task results

Figure 3 shows sensor-space beta- and gamma-band signals recorded during the visuo-motor task. The spatial topographies show average (across all 5 runs) SNR for each sensor, for each frequency band and experimental site. The line plots show the trial-averaged oscillatory envelopes from the sensor with the largest SNR, averaged over the 5 runs with the standard deviation represented by the shaded areas. A time-frequency spectrum for the largest SNR sensor in each case is also shown.

**Figure 3:**
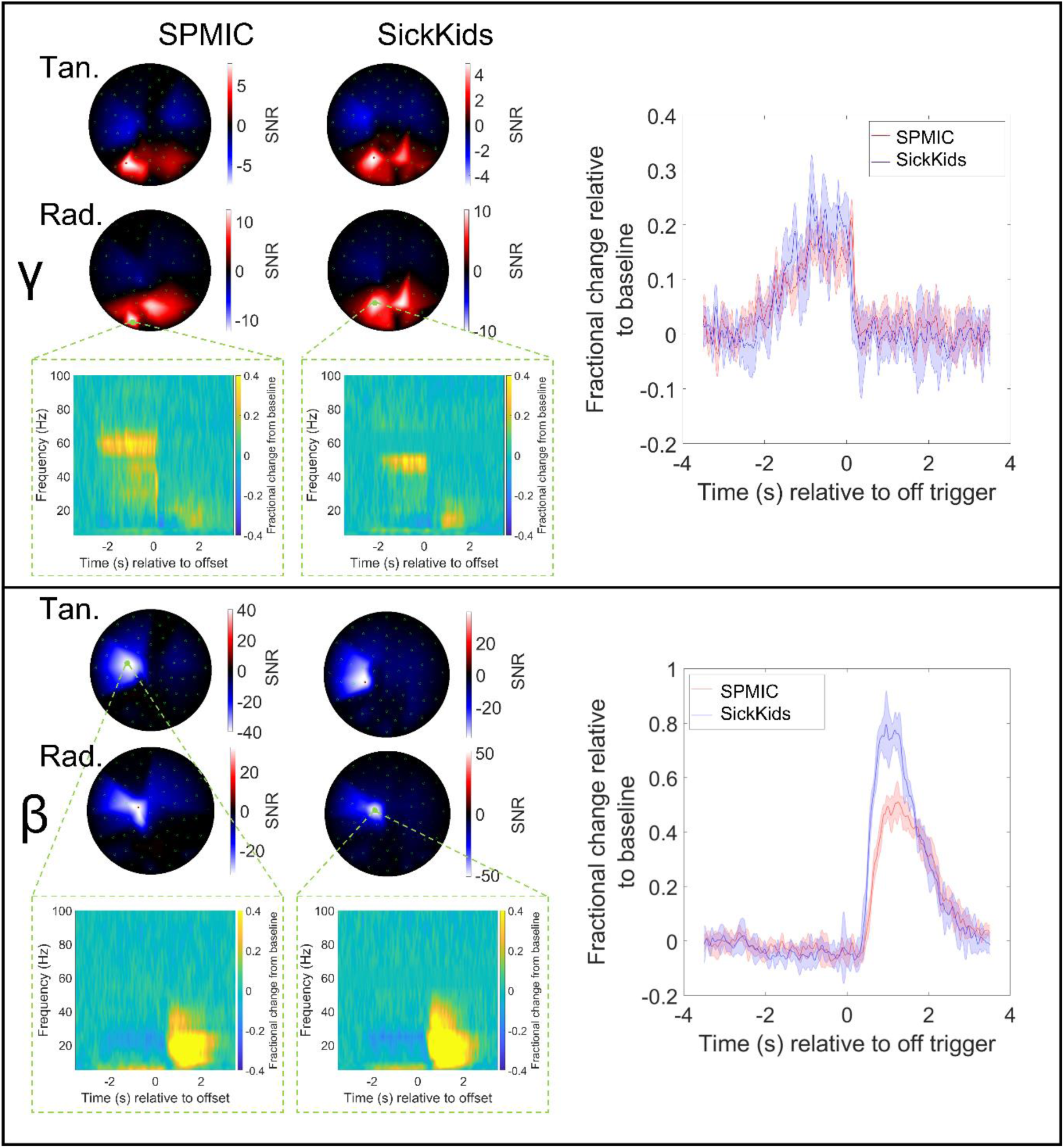
Visuo-motor results (sensor-level). Upper panel: Sensor-level results for the gamma-band (35 – 60 Hz). Spatial topography of the signal-to-noise ratios (SNR) for each sensor averaged across all 5 runs is shown for each site (tangential-axis measurements on top, radial-axis on bottom). On the right, the trial-averaged envelope for the sensor with the highest SNR in the beta band is plotted, with shaded error bars showing the standard deviation across all 5 runs. Results for each site are overlaid, SPMIC in red and SickKids in blue. Time-frequency spectrograms are also shown for the sensor with the highest SNR. Lower panel: Same as the upper panel but in the beta-band (13 – 30 Hz).

In the occipital sensors we observe gamma synchronisation during visual stimulation (in the - 2 s < t < 0 s window). Meanwhile, in the sensors over the motor cortex, we observe the characteristic beta-band desynchronisation (during the −2 s < t < 0 s window), followed by the post-movement beta rebound (during the 0 s < t < 2 s window). While the gamma signal is a similar amplitude for both sites, the beta rebound amplitude is slightly higher for the SickKids site.

Figure 4 shows the results of source reconstruction for the visuo-motor data. The spatial signature of the change in beta and gamma power can be seen for both systems, as well as time-frequency spectrograms (averaged over runs) and the trial-averaged oscillatory envelopes for the peak location of each band. As expected, the beta modulation maps to the contralateral primary sensorimotor cortex, while the gamma modulation maps to primary visual cortex. The Euclidean distance between the SickKids and SPMIC average spatial signature peak locations were 12 mm for the beta peak, and 23 mm for the gamma peak. The source reconstructed time-frequency spectrograms and trial-averaged oscillatory envelopes also show the same characteristic patterns as the sensor-level data with the movement-related beta decrease and post movement rebound, and visual induced gamma amplitude increase both clearly visible; specifically, the temporal correlations between the trial averaged oscillatory envelopes were 0.95 ± 0.01 in the beta band and 0.81 ± 0.05 in the gamma band. The beta-band responses have SNR values of 47 and 48 for SPMIC and SickKids respectively; the gamma-band values are 16 and 8.

**Figure 4:**
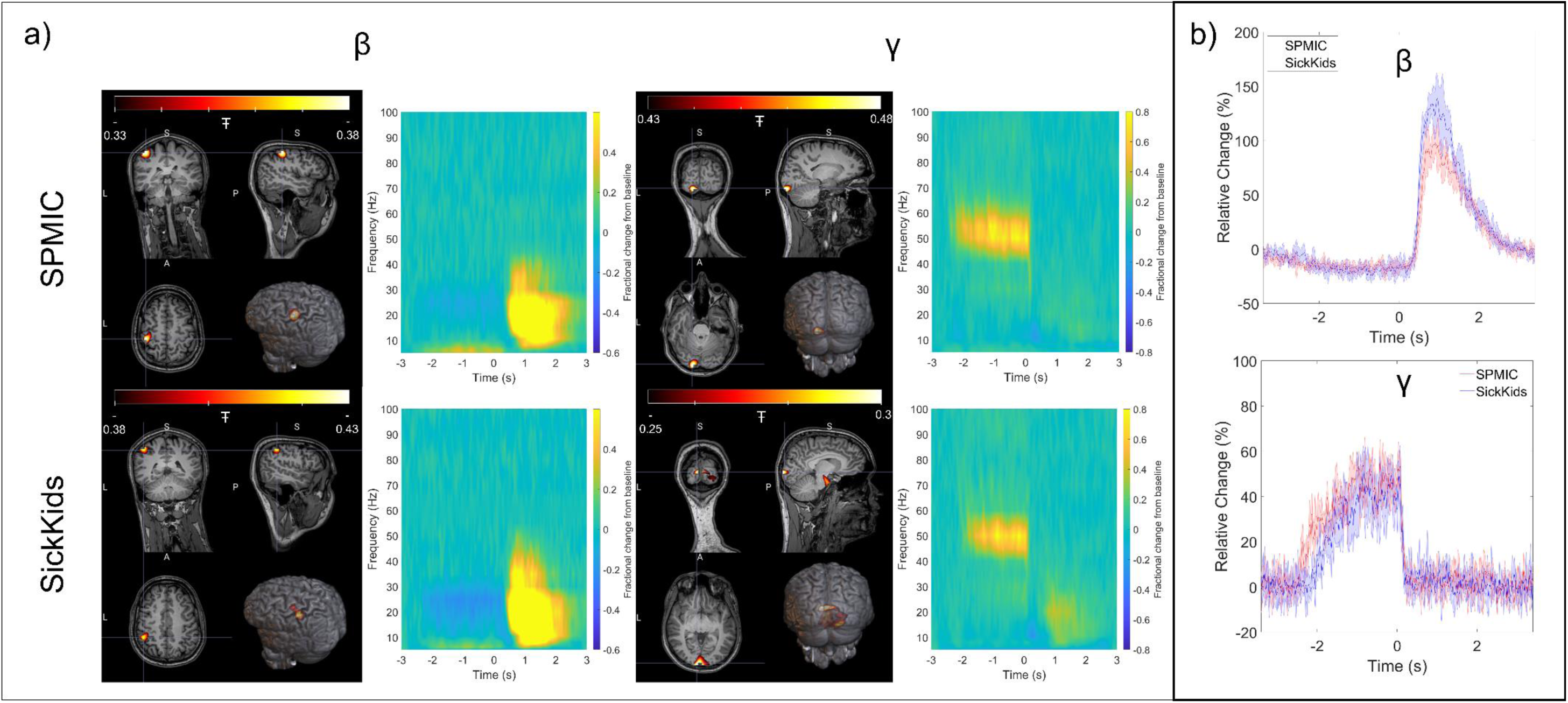
Visuo-motor results (source-level). a) The left and right columns show beta- and gamma-band results respectively. The upper and lower rows show data from the SPMIC and SickKids sites. In each case, a pseudo-t-statistical image, showing the spatial signature of oscillatory modulation (averaged over all 5 runs) is shown on the left, and a time-frequency spectrum for the locations of peak modulation on the right. b) The trial averaged oscillatory envelopes for the beta- (top) and gamma-band (bottom) with SPMIC shown in red, and SickKids in blue with shaded error bars showing the standard deviation across all 5 runs.

### 3.4: Face processing task results

Figure 5 shows a comparison of the evoked responses measured at the sensor level at both sites during the face processing task. The left-hand figure shows the evoked response from the single “best” sensor; SPMIC data shown in red and SickKids data shown in blue. In both cases, data are averaged across all five runs with the shaded area showing the standard deviation across runs. Both sites show similar responses with a peak in field occurring at ~100 ms post stimulation. The field maps on the right of the Figure show the spatial topography of magnetic field across sensors. The two maps show a similar field distribution with a clear dipolar pattern across the occipital sensors in the Z (radial) components. Note the variation between sites is likely because the helmet was positioned differently on the participant’s head.

**Figure 5:**
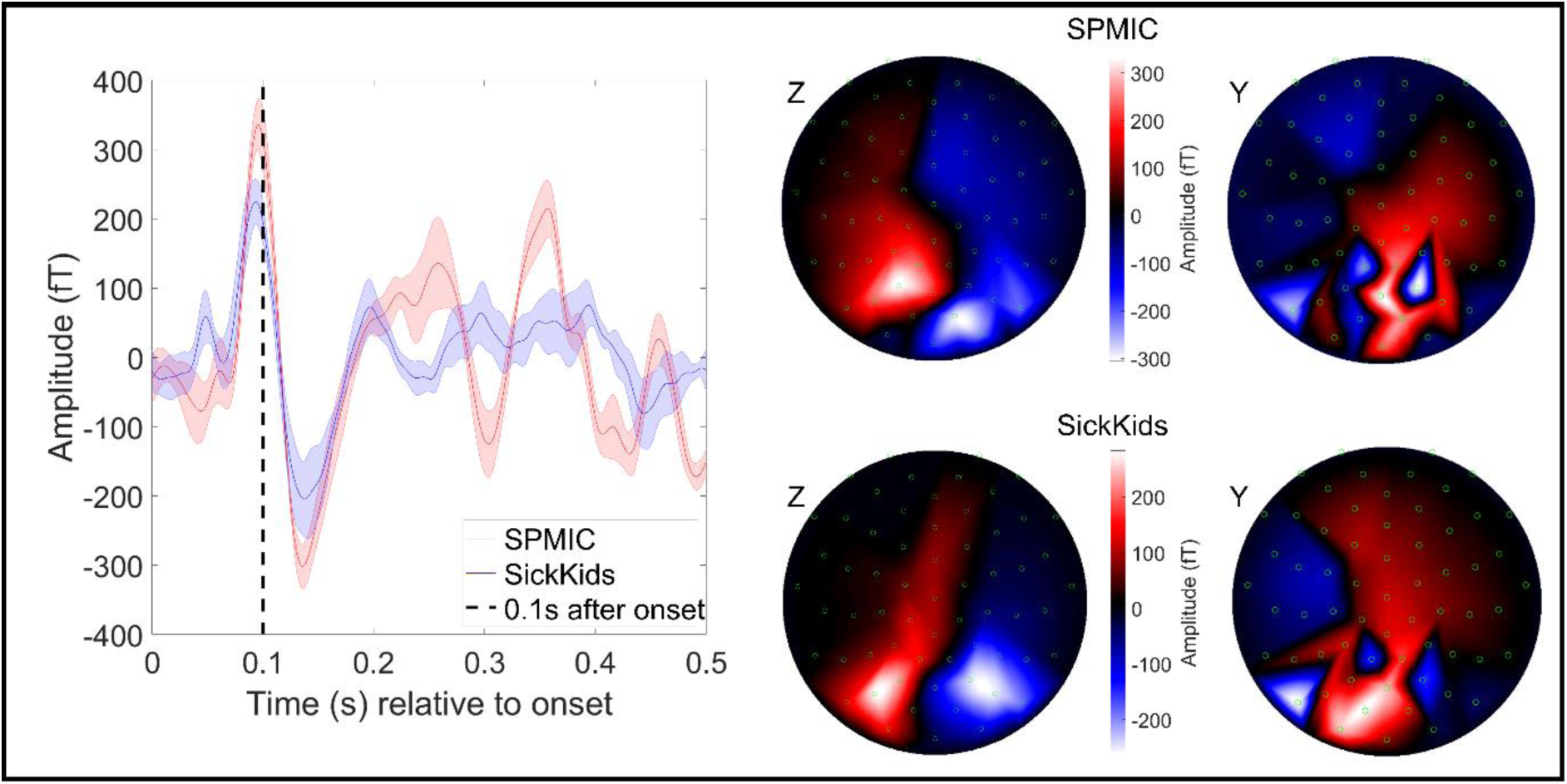
Face processing results (sensor-level). The trial-averaged response in the best sensor for all 5 runs at each site averaged over runs, with the standard deviation across runs shown by the shaded error bars. The best sensor was determined by the range of the trial-averaged signal for each sensor in the 0.075 s < t < 0.175 s window. The field maps on the right show the field distribution at the peak of the average evoked response at 0.1 s (Z-axis (radial) measurements on the left, Y-axis (tangential) on the right).

Finally, Figure 6 shows the source localisation and reconstruction of the evoked response. The peak in the pseudo-t-statistical image is found at the border of the left temporal and occipital lobes for both sites (MNI coordinates: (−46, −68, −11) mm and (−46, −64, −8) mm for SPMIC and SickKids respectively). Two virtual sensor traces are also shown, one extracted from primary visual cortex, and the second from the left fusiform areas for each site (represented as red and blue lines respectively). Inset, the primary visual response is magnified to show the characteristic 75 ms and 145 ms latency peaks. Data from both sites are highly comparable. Spatially, the peak locations in the temporal lobe are separated by 5.7 mm. In terms of temporal morphologies, the two sites are also very similar with near identical latencies for the peak responses, and comparable amplitudes at both virtual sensor locations. Quantitatively, the temporal correlations between the trial averaged evoked responses (measured in the 0 s to 0.3 s window) were 0.70 ± 0.12 in the primary visual area and 0.88 ± 0.04 in the fusiform area. These again show the strong similarity of measured responses across the two sites.

**Figure 6:**
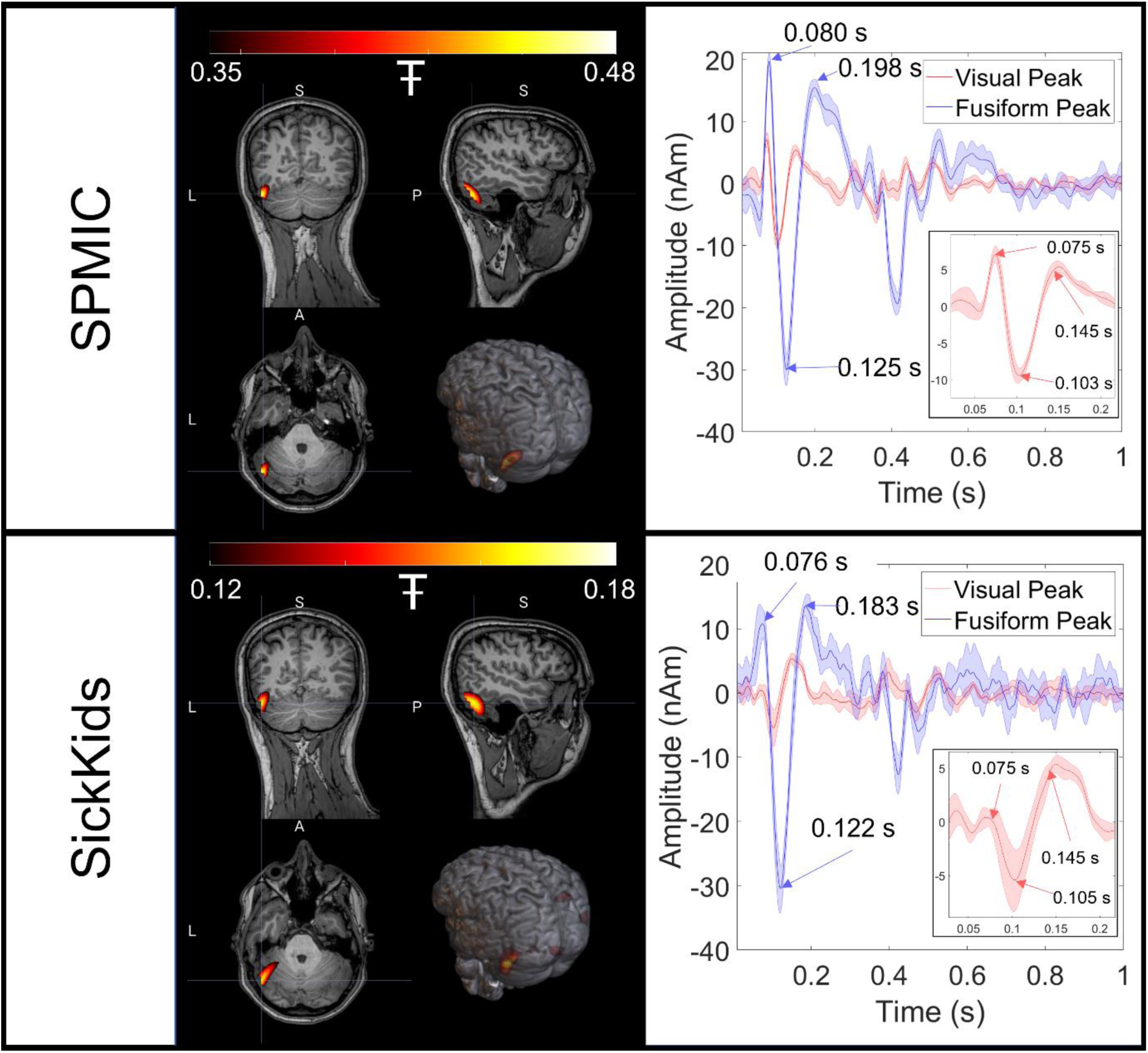
Face processing results (source-level). For each site, the spatial signature of evoked power in the 0.075 s < t < 0.175 s window is shown contrasted against power in the 1.075 s < t < 1.175 s window. On the right, two trial-averaged evoked responses are displayed with the shaded error bars showing the standard deviation across runs. The red line corresponds to a time-course reconstructed in the primary visual area, and the blue line to the left fusiform. Note the differences between regions, but also the similarities across sites. Inset, the primary visual response is magnified.

## 4: DISCUSSION

In this paper, we have shown the first cross-site OPM-MEG comparison. Two near identical systems were constructed in different magnetic environments, with a ~25-fold difference in static background field and a ~30-fold difference in low frequency drift. We showed that, through a combination of background field control (dynamic nulling) and post-processing techniques (homogeneous field correction), OPMs not only remain operational in the non-optimised magnetic environment, but also demonstrate a noise floor of ~16 fT/sqrt(Hz) which is only slightly higher than our low noise environment (~10 fT/sqrt(Hz)). In our human experiments, with the same participant scanned multiple times, we were able to record robust, high-quality MEG data in both environments. Specifically, we were able to reconstruct both beta- and gamma-band modulation in our visuo-motor task, and evoked responses in our face processing paradigm. On average, the spatial discrepancy between localisations at the two sites was of order 10 mm. The temporal correlation between the sites was 0.82 ± 0.06 (collapsed across runs and tasks). Thus, both sites showed highly comparable signals, demonstrating that OPM-MEG systems can work reliably and similar to a low-noise environment, even in busy city centre sites.

Low frequency drift in background field is a significant issue for OPMs, since a shift away from zero has a marked effect on sensor gain. Specifically, a change in field of 3.5 nT causes a change in gain of ~5%; the levels of drift observed in the MSR at the SickKids site were of order 10 nT, which would correspond to ~30% gain change. Without dynamic nulling and field correction, this would be sufficient to invalidate the models used for source reconstruction and render the recorded data an inaccurate representation of brain activity. Previous work has shown that dynamic nulling is effective at cancelling low frequency field drifts (Holmes et al., 2019), although existing demonstrations have been based on lower amplitude artefacts. Here results in Figure 2 show that low frequency drifts up to 10 nT could be controlled successfully, allowing the sensors to remain within their operational range, measuring high fidelity MEG data. Dynamic nulling is therefore critical to enable successful OPM-MEG operation. However, the precise methodology used has some constraints. First, the reference array used included only 4 sensors at both sites; whilst this enables accurate characterisation of background field close to the helmet, field gradients can only be calculated in a single orientation (Z). Second, the power spectra in Figure 2 show that dynamic nulling impacts on interference at higher frequencies. This is due to dynamic range: we have to generate fields of order 10 nT, and the dynamic range of the current drivers means a least significant bit size of ~1 pT. Consequently even the smallest changes in current through the coils generate a shift in background field which is large relative to the target noise floor (15 fT/sqrt(Hz)). Thus, any noise is transmitted to the OPMs around the head by the coils themselves, raising the noise floor at all frequencies. Whilst our dynamic nulling scheme worked well, improved reference array design and lower noise current drivers would likely further improve recordings.

Although interference was high following dynamic nulling, it was adequately controlled via the application of homogeneous field correction, which was able to reduce the noise floor at high frequencies (10 Hz – 40 Hz) from ~100 fT/sqrt(Hz) to ~16 fT/sqrt(Hz). HFC is an attractive solution to removal of interference in an OPM array; it is simple to implement, and the low rank of the model (i.e., the assumption of homogeneity across the head sensors) means a low likelihood of removing neural signal. This is especially important in OPM arrays as they typically contain fewer sensors than a SQUID array, hence the likelihood that any spatial basis set will explain the neural signal by chance is increased. However, HFC is extremely dependent on knowledge of the relative orientations of each sensor. For rigid additively manufactured helmets as used in the current study, where the relative sensor locations and orientations are precisely known (from the electronic CAD file used to define the print), this issue is reduced. However, if flexible (EEG-like) caps were used (e.g., as in Hill et al., 2020) the relative sensor locations and orientations are more challenging to measure and could even change throughout an experiment. It is therefore likely that the utility of HFC would decline dramatically in this case.

Despite limitations of both the interference rejection methods used, we showed that systems at both sites produced high fidelity data. Spatially, we observed excellent agreement in source localisation for both the beta band effects in sensorimotor cortex (~10 mm), and the evoked response in left fusiform area (~5 mm). The spatial differences across sites for the gamma band effects in visual cortex were larger (~28 mm). There are likely several reasons for this. First the stimulus was a centrally presented circular grating, which means the spatial extent of the cortical regions activated will be large. In addition, depending on precisely how the screen was set up, and how the subject viewed it, the retinotopic organisation of the visual cortex is likely to result in a spatial shift of the peak in the response. Finally, gamma is well known to be low amplitude and consequently low signal to noise response. These effects combined will likely cause the spatial difference to be higher than for the motor response (which will be mapped to the motortopic representation of the same finger) and the fusiform. Temporally, all responses showed good agreement, with beta, gamma, and fusiform evoked responses all exhibiting >80% correlation. In addition, all responses at both sites were of comparable amplitude (the slight discrepancy in the beta rebound amplitude at the sensor level is likely due to sensor placement; this was largely rectified by source reconstruction).

A limitation of the SickKids site remains the ambient field. Even after dynamic nulling and HFC are applied, sources of interference remain due to the busy clinical environment and city centre infrastructure in which the system is located. Such interference, which is likely due to e.g., vibrations in the MSR (for example, from a car driving underneath) led to a greater number of trials being rejected at the SickKids site, compared to the SPMIC site (particularly in the face processing paradigm). Despite this, the system is still fully operational, and a simple trial rejection algorithm was able to discard trials with high degrees of interference. In future installations, the use of an OPM-optimised magnetically shielded rooms, with the capability to demagnetise the inner mu-metal walls (using a degaussing system) will likely reduce these effects.

## CONCLUSION

We have demonstrated that a commercial OPM-MEG system can be sited in a non-optimised shielded room, in a clinical setting, in a major city and yield data comparable to that collected in an optimised site. Through use of dynamic nulling and homogeneous field correction, this system exhibits low noise, and successfully records OPM-MEG data which are well matched to equivalent data collected using existing (“tried and tested”) OPM-MEG instrumentation. We stress that the system remains nascent technology; further work can be done on artefact rejection, with more sophisticated post-processing techniques being explored. In addition, both OPM systems will benefit from a larger sensor array, which will improve coverage of the brain, and increase the information available to algorithms like HFC and source localisation, which will further improve signal to noise ratio. Despite this, the paper shows that “plug-and-play” OPM-MEG systems now exist, they can be easily sited even in challenging environments, generate high fidelity data, and will provide significant benefit to clinical research groups who can now begin to exploit the high degrees of practicality and lifetime compliance which OPMs afford, to explore clinical questions.

## ACKNOWLEDGEMENTS

We acknowledge support from the UK Quantum Technology Hub in Sensing and Timing, funded by EPSRC (EP/T001046/1). Sensor development was made possible by funding from the National Institutes of Health (R44MH110288). We also express thanks to Metamorphic AM and Added Scientific Ltd. for the useful and productive discussions that led to the design of the lightweight helmet used for paediatric measurements.

SickKids also acknowledges the following grants:

- Brain oscillations and emerging symptoms in toddlers with autism: Combined insights from new MEG technology and basic neurophysiology.. Safar, K., Anagnostou, E., Brian, J., Collingridge, G., Taylor, M.J. Feiga Bresver Catalyst Grant. April 1, 2021 – March 31, 2023.
- It’s all about time: Optimising infrastructure for functional brain imaging in children. Taylor, M.J., Mabbott, D.J., Dunkley, B.T. CFI-JELF, 2018-2023,

